# Comparative analysis of the effects of retinoic acid versus paclitaxel and everolimus on HL60 cells proliferation and viability

**DOI:** 10.1101/2023.04.03.535342

**Authors:** Athanasia Zampouka, Triantafyllia Papadimitropoulou, Maria Salagianni, Maria Vaiou, Amalia I Moula, Athanasios Giannoukas, Anargyros N Moulas

## Abstract

**Purpose:** All trans-retinoic acid (atRA) has been proposed as a novel drug for drug eluting stents (DES). Currently complications of DES have been at least partially attributed to the drugs that are used: paclitaxel and sirolimus and its derivatives like everolimus. We compared the effects of atRA, paclitaxel and everolimus on the proliferation and viability of human leukemia cells (HL60).

**Methods:** Cells were cultured with 0.1μM and 10μM of atRA, paclitaxel or everolimus. Cell proliferation and viability was evaluated with trypan blue at 24, 48 and 72 hours.

**Results:** All drugs caused a statistically significant, dose-dependent reduction of cell proliferation rate from the first 24 hours. atRA and everolimus did not affect cell viability as the treated cells showed high viability (95-98%), while paclitaxel decreased significantly the viability to below 16% at 72 hours. Unlike the cytotoxic effect of paclitaxel on HL60, atRA demonstrated a cytostatic effect comparable to everolimus.

**Conclusion:** The ability of atRA to limit cell proliferation without affecting cell viability in a manner similar to everolimus, highlights its potential to be used on DES as a novel drug for treatment of restenosis with potentially minimal side-effects. Further research with different cell types, is needed in order to elucidate the possible usefulness of RA on DES.

## 1. Introduction

Paclitaxel and rapamycin (sirolimus) and its derivatives such as everolimus are currently used on endovascular peripheral and coronary drug eluting stents (DES) for treatment of restenosis[1]. Paclitaxel acts by binding with microtubules, leading to arrest of the cell cycle and ultimately causing cell death[2]. Rapamycin and its derivatives act by inhibiting the mTOR (mammalian target of rapamycin), a protein that regulates the proliferation, growth and survival of cells[3]. Latest generation DES are quite safe and effective, however they still harbor risks of potentially lethal complications, such as late stenosis and thrombosis. As these effects have been at least partially attributed to the toxicity of the drugs currently in use, research is ongoing on new less toxic candidate drugs.

Retinoic acid (RA) is a naturally occurring vitamin A (retinol) derivative with important biological actions including regulation of cell differentiation and proliferation. The all trans-isomer of RA (atRA) is used as treatment for disorders related to cell proliferation, including acne and acute myeloid leukemia. RA acts mainly by binding to specific nuclear receptors (namely retinoic acid receptors or RAR and retinoid X receptors or RXR) and affecting the expression of specific genes[4].

RA has been reported in a number of *in vitro*[5] and *in vivo* studies[6-8] to reduce neointimal proliferation and it has been proposed as an alternative drug for DES[9], based on its potential efficacy to minimize the side effects of the drugs currently use in DES. Recently, RA eluting stents were found not to be related with thrombus formation and having an acceptable degree of stenosis in a rabbit iliac artery model[9].

Herein, for the first time we sought to compare the effects of RA on cell proliferation with drugs already used on commercial stents, particularly paclitaxel and everolimus. For the purpose of this study, we used the HL60 suspension cell line of promyeloblasts derived from a 36-year old woman with acute promyelocytic leukemia. This cell line has a stable proliferation profile which was ideal for this comparative study of the efficacy of paclitaxel, everolimus and atRA with respect to cell viability and proliferation.

## 2. Materials and Methods

### 2.1. Cell culture

Cell culture Human Leukemia cell line HL60 was purchased from the European Collection of Authenticated Cell Cultures (ECCAC, Health Protection Agency, Salisbury, UK). The cells were cultured in RPMI 1640 (Gibco) medium with 1% L-Glutamine 200 mM (Gibco), 1% Penicillin/Streptomycin Solution 100x (Biosera) and 20% of heat-inactivated FBS (Gibco) in incubator at 37°C, with 5% CO_2_ until they reached log phase. In standard culture conditions, the FBS was reduced to 10% and the cells were passaged every 2 days at a concentration of 0.5 × 10⁶ cells/mL. All experiments were performed using logarithmically growing cells.

### 2.2. Drugs and chemicals

Absolute ethanol was purchased from Panreac Applichem. All-trans retinoic acid, Everolimus and PTX were purchased from Cayman Chemical (Michigan, USA), dissolved in absolute ethanol to a stock solution of 10 mM, and sterile-filtered. All preparations were performed under light protection for atRA and prepared freshly prior to each application. Two final concentrations of each drug in the incubation solution were tested, 0.1 μmol/l (μM) and 10 μmol/l.

### 2.4. Drug assay on cells

The experiments were performed in 12-well plates (Corning®) with initial cell concentration of 10⁵ cells/well at zero time (T_0_). The concentrations of 0.1 μΜ and 10 μΜ of atRA, everolimus and paclitaxel were tested in duplicates in each experiment for the selected time points of 24, 48 and 72 hours. Cells treated with vehicle (0.1% ethanol) were used as a negative control. All experiments were conducted in triplicate.

### 2.4. Trypan blue assay

The assessment of drug effect on cell proliferation and viability was done with the trypan blue assay. On each time point, the cell suspension from every well was collected in a 1.5 mL tube individually and a sample of it was diluted with Trypan Blue Solution 0.4% (Gibco) and viewed under light microscope (Zeiss) using a Neubauer chamber. The cell proliferation was expressed as total count of viable cells per condition using the formula:

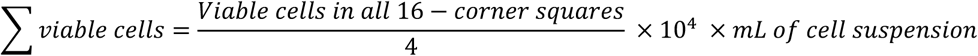

The % cell viability was calculated using the formula below:

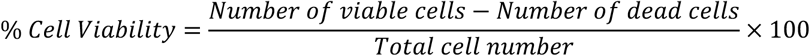

### 2.4. Statistics

The SPSS Statistical package for Windows (IBM Corp.) was used. Data are expressed as mean ± standard deviation (SD) from three independent experiments. Student’s t-Test was used for the comparisons of means and the statistical significance limit was set at p < 0.05.

## 3. Results

### 3.1. Cell proliferation

The effect of atRA, Everolimus and paclitaxel on cell proliferation was examined by counting the cells at 24, 48 and 72 hours after the addition of the respective pharmacological agents. Treatment of HL60 with the above mentioned drugs resulted in statistically significant reduction of cell proliferation rate from the first 24 hours in a dose dependent manner (Figure 1A). Paclitaxel showed the most prominent effect, resulting in 34.6% and 26.8% reduction of proliferation at 24 hours for 0.1μM and 10μM respectively, reaching an almost complete growth inhibition at the longer incubation times. Everolimus at 0.1μM concentration resulted in 85.9% reduction of proliferation at 24 hours, 80.4% reduction at 48 hours and in 70% reduction at 72 hours. A more prominent effect was achieved upon exposure to 10 μM everolimus, resulting in a statistically significant attenuation from 80.6% at 24 hours to 55.5% at 72 hours. (Figure 1A, 1B & 1C). atRA also reduced the proliferation from 87.7% at 24 hours to 75.9% at 72 hours for 0.1μM, although in this concentration, its effect at 48 hours was not statistically significant. At the high concentration (10μM), atRA excibited a more pronounced anti-proliferative effect similar to that observed for everolimus but still lower of that of PTX.

**Figure 1:**
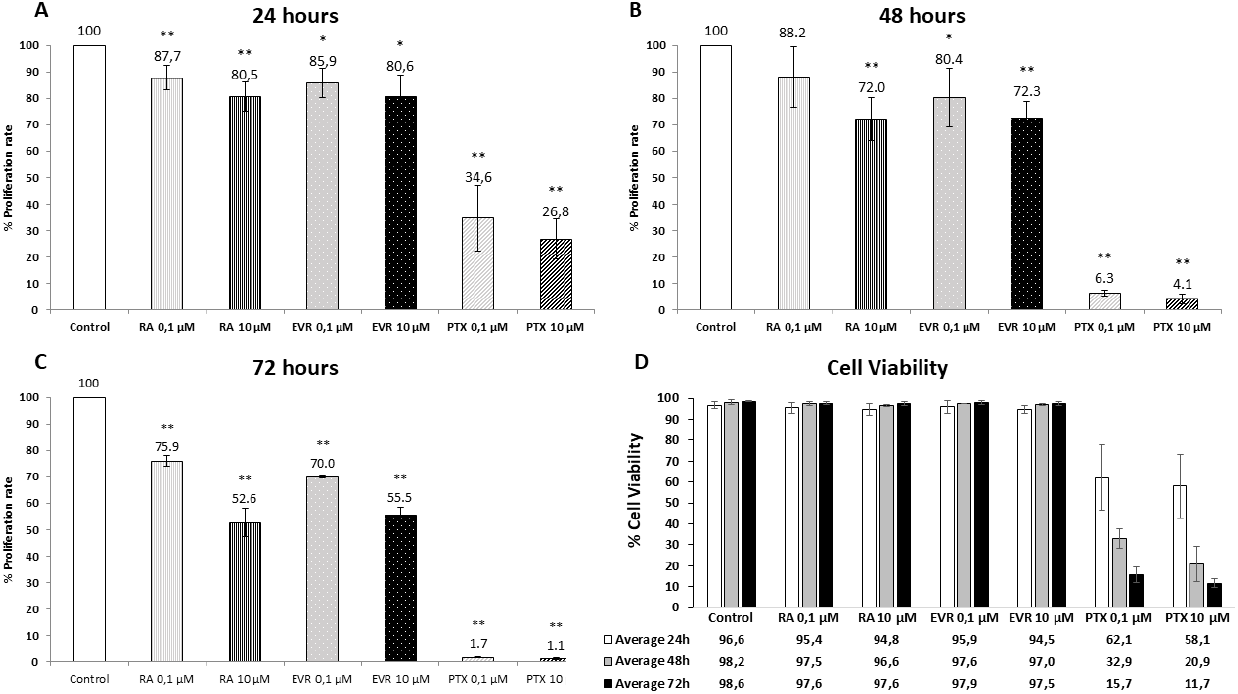
The effect of atRA, Everolimus and Paclitaxel on cell proliferation and cell viability. A) The % proliferation rate of HL60 cells with 0.1 μM and 10 μM concentrations of atRA (RA), everolimus (EVR) and paclitaxel (PTX) treatment after 24, B) 48 and C) 72 hours. D) The % cell viability of HL60 cells with 0.1 μM and 10 μM concentrations of drugs after 24, 48 and 72 hours. *P<0.05, **P<0.01 compared to control, Student’s t-test. The data were derived from 3 independent experiments.

### 3.2. Cell viability

To assess the possible cytotoxic effects of these drugs, we proceeded to inspect HL-60 for evidence of viability changes by Trypan blue labeling of cells. atRA and everolimus, even at 10uM, did not promote changes in HL-60 viability over a period of 72 hours and the treated cells showed viability percentages in the range 94.8-97.9%, with no significant differences to control condition (range 96.6-98.6) for the 3 time points. (Figure 1D). In contrast to the other two drugs, paclitaxel caused a statistically significant decrease of the viability of cells averaging 62.1% for the0.1 μΜ and 58.1% for the 10 μΜ at 24 hours and reaching 15.7% and 11.7% respectively at 72 hours (Figure 1D).

## 4. Discussion

atRA was compared with paclitaxel and everolimus, two drugs with different mechanisms of action that are already in use in DES, for their effects on cell proliferation, the main key contributor to vessel restenosis.

atRA significantly slowed down the proliferation of HL-60 cells in a dose-dependent manner over the course of 72 hours but did not affect their viability levels. These findings suggest that atRA-induced decline in HL-60 proliferation is due to a pharmacological effect on cellular signaling phenomena that control cell division, rather than cytotoxic effects of this drug. Previous studies support these findings as RA is known to induce HL-60 differentiation and arrest cell growth at G1/0 phase of cell cycle[10, 11]. Paclitaxel had a cytotoxic effect on the cells by significantly reducing their proliferation ability and viability. Paclitaxel studies on HL-60 cells demonstrated similar apoptotic effects and cell growth arrest, in tests of the drug for the same time points[12-14]. Lastly, everolimus caused a significant but moderate reduction on leukemia cell proliferation compared to paclitaxel without significantly affecting their viability and this observation is in accordance with previous studies[15]. Unlike the cytotoxic effect of paclitaxel on HL-60, atRA acted in a cytostatic fashion similar to that observed for everolimus.

Our results indicate that the effect of atRA is less prominent in the lower tested dose of 0.1 μM, compared to paclitaxel and everolimus. However, at the 10μM dose, atRA achieved a high degree of proliferation inhibition comparable to that of everolimus but still lower of that of paclitaxel. These observations suggest that for possible applications on drug eluting stents, the necessary dose of atRA should probably be higher than the respective dose of paclitaxel and everolimus by a factor of 10 times or more. Also longer incubation times seem to improve further the anti-proliferative effect of the atRA in a similar manner observed for PTX and Everolimus.

As previously mentioned, paclitaxel and rapamycin derivatives are used in currently available DES. Their physical properties, such as in vivo half-life and lipophilicity, as far as their anti-restenotic efficiency supported their prolonged use in the clinic[16]. However, several meta-analyses comparing long-term effects of paclitaxel-eluting and everolimus-eluting stents after implantation in patients showed that although DES with everolimus appeared to be overall safer and more effective, there were still clinical phenomena such as stent thrombosis, late luminal loss, target lesion and target vessel revascularization[17-20]. Therefore, the need for new drugs that limit the long-term, adverse effects of DES in combination with the preliminary results of this comparative study call attention to further research on retinoic acid’s potential use on DES.

RA is a naturally occurring compound with relatively low toxicity. Its ability to limit cell proliferation without affecting cell viability in a way similar to everolimus, highlights its potential to be used on DES as a novel drug for treatment of restenosis with potentially minimal side-effects. Further research, including different cell types and *in vivo* experiments, is needed in order to elucidate the role and possible usefulness of RA on DES for the treatment of restenosis.

## Statements and Declarations

### Availability of data and material (data transparency)

The manuscript has no associated data.

### Code availability (software application or custom code)

Not applicable.

### Author Contributions

All authors contributed to the study conception and design. Material preparation, data collection and analysis were performed by Athanasia Zampouka, Triantafyllia Papadimitropoulou, Maria Salagianni, Maria Vaiou and Amalia I Moula. The first draft of the manuscript was written by Athanasia Zampouka and Anargyros Moulas and reviewed by Athanasios Giannoukas and all authors commented on previous versions of the manuscript. All authors read and approved the final manuscript.

### Ethics approval

Not applicable.

### Consent to participate

Not applicable.

### Consent for publication

Not applicable.

## Acknowledgement

We would like to thank Evangelos Andreakos for helpful suggestions and discussions.

